# Neural adaptation to accented speech: prefrontal cortex aids attunement in auditory cortices

**DOI:** 10.1101/852616

**Authors:** Esti Blanco-Elorrieta, Laura Gwilliams, Alec Marantz, Liina Pylkkänen

## Abstract

Speech is a complex and ambiguous acoustic signal that varies significantly within and across speakers. A prevalent and ubiquitous example of such variation is accented speech, to which humans adapt extremely rapidly. The goal of this study is to uncover the neurobiological bases of the attunement process that enables such fluent comprehension. Twenty-four native English participants listened to words spoken by an unaccented “canonical” American talker and two “accented” talkers, and performed a word-picture matching task, while magnetoencephalography (MEG) was recorded. Accented speech was created by including systematic phonological substitutions within the word (e.g. [s] → [sh]). Activity in the auditory cortex (superior temporal gyrus) was greater for accented speech, but, critically, this was not attenuated by exposure. By contrast, prefrontal regions showed an interaction between the presence of an accent and amount of exposure: while activity decreased for canonical speech over time, responses to accented speech remained consistently elevated. Grainger causality analyses further revealed that prefrontal responses serve to modulate activity in auditory regions, suggesting the recruitment of top-down processing to decode accented signal. In sum, our results show that accented speech does not elicit the same prefrontal reduction in amplitude over time that unaccented speech does, and points to a dynamic exchange of information between the prefrontal and auditory cortices in order to recalculate phonetic classification and subsequent identification of lexical items.

**Significance statement:** Human ability to adapt to different people’s idiosyncratic pronunciations is a hallmark of speech comprehension. This study aims to address whether adaptation to speakers’ accents manifests itself at the perceptual level (i.e. through adaptation of low-level neural responses in the auditory cortex) or at the post-perceptual level (i.e. higher-order regions correcting the sensory signal received from auditory cortex). Our results support the post-perceptual hypothesis: we found that responses in auditory cortex emitted an error-like signal that was invariant to exposure; whereas responses in the prefrontal cortex were modulated by exposure. These findings provide initial insight into accent adaptation, and illuminate the computational stages supporting speech comprehension more generally.

## INTRODUCTION

Speech provides the ability to express thoughts and ideas through the articulation of sounds. Despite the fact that these sounds are often ambiguous, humans are able to decode these signals and communicate efficiently.

This ambiguity emerges from a complex combination of physiological and cultural features (Babel & Munson, 2014; Baken & Orlikoff, 2000; Benzeghiba et al., 2007; Labov, 1972; Nolan, 1980; Pierrehumbert, 2003); and results in situations where, for example, one talker’s “ship” is physically very similar to another speaker’s “sip” (Newman, Clouse & Burnham, 2001). Indeed, any given sound has a potentially infinite number of acoustic realizations (Allen, Miller & DeSteno, 2003; Kleinschmidt & Jaeger, 2015); comprehending speech therefore necessitates accommodating for the discrepancy between the perceived phoneme and the personal ideal form of that phoneme. This process is known as perceptual attunement (Aslin & Pisoni, 1980).

Accommodation is particularly critical for comprehension when listening to an accented talker (Mattys et al., 2012), and involves recalibrating the mapping from acoustic input to phonemic categories through perceptual learning (Aslin & Pisoni, 1980; Norris et al., 2003; Kraljic and Samuel, 2005, 2006, 2007; Maye et al., 2008; see Samuel & Kraljic, 2009 and Guediche et al., 2014 for a review). This adjustment happens quickly: within the first 10 sentences for non-native speech (Dupoux & Green, 1997; Clarke & Garrett, 2004), and within 30 sentences for noise-vocoded speech (Davis et al., 2005).

Although perceptual adaptation in speech comprehension has been robustly reported, the neural underpinnings supporting this process remain poorly described. So far, it is only known that the patterns of activation underlying the perception of speech in noise and accented speech dissociate (Adank et al., 2012), and that they vary when listening to regional as compared to foreign accents (Goslin et al., 2012). However, a lot is known about how *canonically* pronounced phonemes are processed. A collection of neuro-physiological studies have linked responses in superior temporal gyrus (STG) at 100 ms latency with phonetic feature processing (Mesgarani et al., 2014; Di Liberto, 2015), which critically maps to the perceived phonological category and is not modulated by within-category variance (Chang et al., 2010). This processing stage may therefore be linked to the perceptual stage of speech processing. Neural responses later than 100 ms appear to be associated with post-perceptual processing, such as the integration of previous expectations (e.g., phoneme surprisal; Ettinger, Linzen & Marantz, 2014; Gwilliams & Marantz, 2015) and integration of lexical information (Gow, 2008).

As the first study to investigate the neural processes supporting accented speaker adaptation, we start with a basic but fundamental question: Does the re-mapping between acoustic input and phonological percept occur: i) at the perceptual stage of processing (sensory hypothesis), or ii) as a higher-level post-perceptual stage (repair hypothesis)? To address this question, 24 native English participants listened to isolated words that were either pronounced canonically (baseline condition), or were pronounced with systematic phonological substitutions. Neural responses were recorded using magneto-encephalography (MEG), while participants performed a word-picture matching task. We would expect the behavioural correlate of accent adaptation (improved comprehension as a function of exposure) to be mirrored in neural responses in STG at 100 ms if the process affects perception directly; however, we would expect it in frontal areas at a later stage of processing if it reflects a higher-level repair process.

## METHODS

### Stimuli Selection

In order for the experiment to be informative of real-world accent processing, we selected three phonemic minimal-pairs that are often transposed in certain variations of English (Attested variation: v → b, θ → f and ∫ → s substitutions). Importantly, these phoneme substitutions are systematic for speakers of those variations and occur uni-directionally (e.g., native Spanish speakers of English often substitute instances of /v/ with /b/, but not vice versa). We also presented participants with substitutions in the reverse direction, which are not naturally occurring and thus, participants could not have been previously exposed to (Unattested variation; b → v, f → θ and s → ∫). This procedure allowed us to test naturally occurring substitutions while also creating a condition where attunement before the experiment was null. Using the same phonemes in both conditions additionally allowed us to keep the perceptual distance in the two conditions constant, thus ensuring that any potential difference between the two was due to the level of initial attunement and not this confounding factor. Lastly, our experiment included a baseline condition where all phonemes were pronounced accurately and no adaptation was required (Baseline condition; for examples of stimuli for each condition please see Fig. 2A).

**Figure 1.**
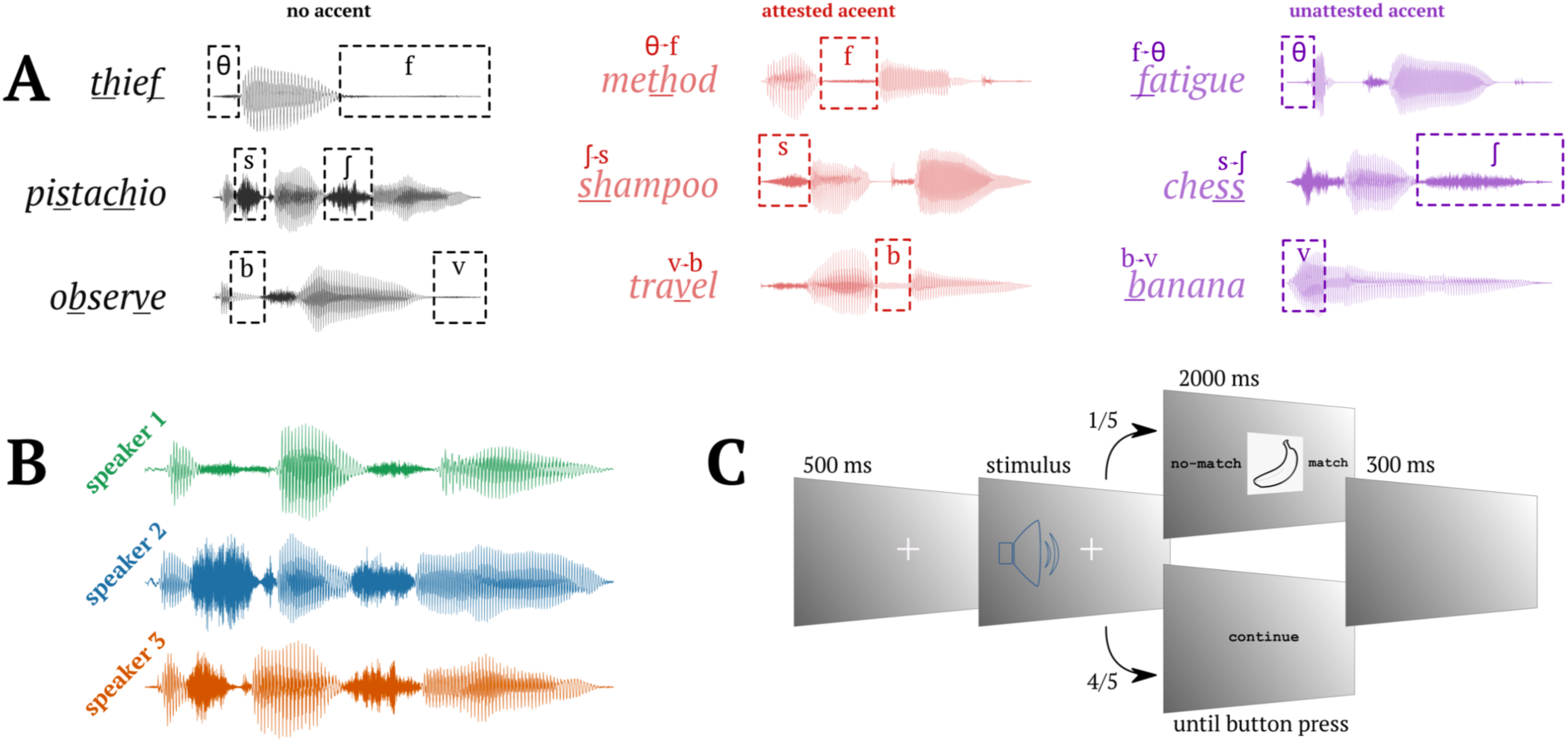
Experimental stimuli and design. (A) Experimental conditions. In black, left column: baseline condition. In red, middle column: attested substitutions. In purple, right column: unattested substitutions. (B) Waveform of three speakers’ pronounciation of *pistaccio*. (C) Trial structure.

**Figure 2.**
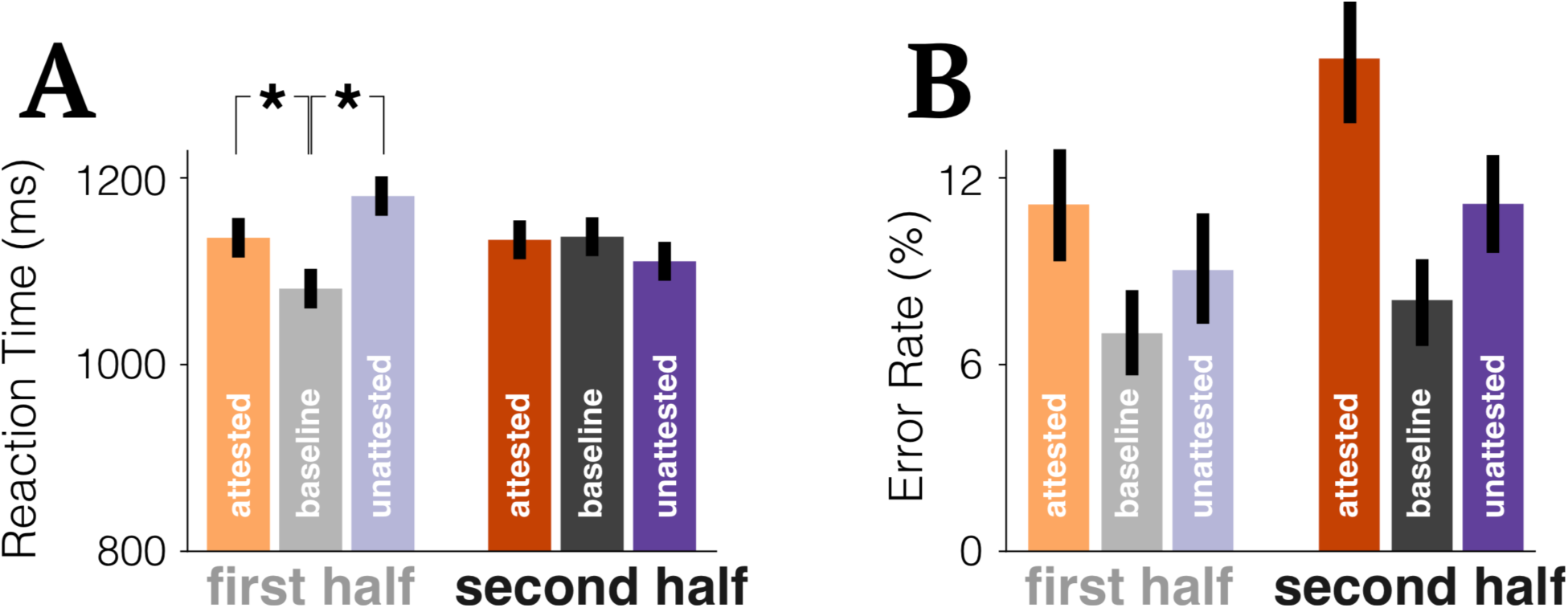
Behavioral responses to the word-picture matching task. A) Behavioral reaction times showing that people are slower identifying pictures at the beginning of the experiment when the word was accented, but adjust to it as the experiment goes on. B) Error rates showing that participants made more mistakes for accented speech – regardless of exposure.

Critical phonemes were presented embedded in meaningful words, and all words were selected such that the critical phoneme substitution created non-words (e.g., the word *travel* becomes a non-word if critical phoneme /v/ is substituted by /b/ to form *trabel*). There were two reasons for selecting these types of words: i) to avoid cases where not adapting to the phoneme substitutions would still result in a meaningful word (and thus the substitution would go unnoticed) and ii) to take advantage of the Ganong effect (Ganong, 1980), which establishes that humans tend to make phonetic categorizations that make a word of the heard strings. It follows then that using cases where only one of the interpretations results in a meaningful word should maximize the chances of listeners’ adaptation.

All stimulus words had a minimum log surface frequency of 2.5 in the English Lexicon Project corpus (Balota et al., 2007), were between 3 and 10 phonemes long, and were monomorphemic. Words across conditions were matched for log surface frequency (M = 8.19, SD = 1.92), length (M = 4.51, SD = 1.36) and critical phoneme position (M = 1.64, SD = 1.36). All critical phonemes appeared the same number of times in the experiment.

### Experimental Design

The experiment comprised of a total of 600 trials, each consisting of the auditory presentation of a single word. Each experimental condition contained 240 critical phonemes. In both variation conditions (Attested and Unattested), each word contained only one critical phoneme substitution, as having multiple substitutions in a single token may impair listeners’ comprehension. This selection exhausted the pool of English words that met all of our stimuli selection criteria. Thus, in the Baseline condition each word instead contained two critical phonemes (120 words, 240 critical phonemes). This did not pose any methodological concern though, as all analyses of MEG data were time-locked to phoneme onset and not to word onset; thus, this paradigm allowed for 240 data points to be collected in the Baseline condition just like in the other two conditions (for a complete list of experimental items please see Additional Materials 1).

Three native speakers of American English were recorded to create the stimuli. Each individual was recorded saying all 600 items, following the variation rules previously explained for the items in the Attested and Unattested conditions, and naturally pronouncing the Baseline words. Stimuli were recorded in a single recording session for each speaker. Each word was read three times, and the second production of the word was always selected to allow for consistent intonation across stimuli. Critical phoneme onsets were identified manually and marked using Praat software (Boersma & Weenink, 2009).

The assignment of speaker to condition was shuffled across participants following a latin-square design. Items for each condition were further divided into blocks of 20 items to form experimental blocks. Items within blocks were fully randomized, and blocks within the experiment were pseudo randomized following one constraint: Two blocks of the same condition never appeared successively. Each stimulus was presented once during the experiment and each participant was presented with a unique order of items. Stimuli were presented binaurally to participants through tube earphones (Aero Technologies), using Presentation stimulus delivery software (Neurobehavioral Systems).

### Experimental procedure

Each trial began with the visual presentation of a white fixation cross on a gray background (500 ms), which was followed by the presentation of the auditory stimulus. In one fifth of the trials, 500 ms after the offset of the auditory stimulus, a picture appeared on the screen. Participants were given 2000 ms to make a judgment via button press to indicate whether the picture matched the word they had just heard. For all participants, the left button indicated “no-match” and the right button indicated “match”. In the other four fifths of the trials, participants would see the word “continue” instead of a picture, which remained on-screen until they pressed either button to continue to the next trial. The picture-matching task was selected to keep participants engaged and accessing the lexical meaning of the stimuli while not explicitly drawing attention to the phoneme substitutions. The 500 ms lapse between word offset and the visual stimulus presentation was included to ensure that activity in our time-windows of interest was not interrupted by a visual response. In all trials, the button press was followed by 300 ms of blank screen before the next trial began (Fig. 2B). Behavioral reaction times were measured from the presentation of the picture matching task.

### Data acquisition and preprocessing

Before recording, each subject’s head shape was digitized using a Polhemus dual source handheld FastSCAN laser scanner (Polhemus, VT, USA). Digital fiducial points were recorded at five points on the individual’s head: The nasion, anterior of the left and right auditory canal, and three points on the forehead. Marker coils were placed at the same five positions in order to localize that person’s skull relative to the MEG sensors. The measurements of these marker coils were recorded both immediately prior and immediately after the experiment in order to correct for movement during the recording. MEG data were collected in the Neuroscience of Language Lab in NYU Abu Dhabi using a whole-head 208 channel axial gradiometer system (Kanazawa Institute of Technology, Kanazawa, Japan) as subjects lay in a dimly lit, magnetically shielded room.

MEG data were recorded at 1000Hz (200Hz low-pass filter), and noise reduced by exploiting eight magnetometer reference channels located away from the participants’ heads via the Continuously Adjusted Least-Squares Method (Adachi et al., 2001), in the MEG Laboratory software (Yokogawa Electric Corporation and Eagle Technology Corporation, Tokyo, Japan). The noise-reduced MEG recording, the digitized head-shape and the sensor locations were then imported into MNE-Python (Gramfort et al., 2014). Data were epoched from 200ms before to 600ms after critical phoneme onset. Individual epochs were automatically rejected if any sensor value after noise reduction exceeded 2500 fT/cm at any time. Additionally, trials corresponding to behavioral errors were also excluded from further analyses.

Neuromagnetic data were coregistered with the FreeSurfer average brain (CorTechs Labs Inc., La Jolla, CA and MGH/HMS/MIT Athinoula A. Martinos Center for Biomedical Imaging, Charleston, MA), by scaling the size of the average brain to fit the participant’s head-shape, aligning the fiducial points, and conducting final manual adjustments to minimize the difference between the headshape and the FreeSurfer average skull. Next, an ico-4 source space was created, consisting of 2562 potential electrical sources per hemisphere. At each source, activity was computed for the forward solution with the Boundary Element Model (BEM) method, which provides an estimate of each MEG sensor’s magnetic field in response to a current dipole at that source. Epochs were baseline corrected with the pre-target interval [-200 ms, 0 ms] and low pass filtered at 40 Hz. The inverse solution was computed from the forward solution and the grand average activity across all trials, which determines the most likely distribution of neural activity. The resulting minimum norm estimates of neural activity (Hämäläinen & Ilmoniemi, 1994) were transformed into normalized estimates of noise at each spatial location, obtaining statistical parametric maps (SPMs), which provide information about the statistical reliability of the estimated signal at each location in the map with millisecond accuracy. Then, those SPMs were converted to dynamic maps (dSPM). In order to quantify the spatial resolution of these maps, the pointspread function for different locations on the cortical surface was computed, which reflects the spatial blurring of the true activity patterns in the spatiotemporal maps, thus obtaining estimates of brain electrical activity with the best possible spatial and temporal accuracy (Dale et al., 2000). The inverse solution was applied to each trial at every source, for each millisecond defined in the epoch, employing a fixed orientation of the dipole current that estimates the source normal to the cortical surface and retains dipole orientation.

### Statistical Analysis

#### ROI analysis

Based on previous research, we focused our analysis on auditory cortex and frontal cortex (Di Liberto, 2015; Mesgarani et al., 2014; Van Engen & Peelle, 2014). We used *freesurfer* to parcellate the average brain (using the *aparc* parcellation) and to index the vertices corresponding to transverse temporal gyrus, superior temporal gyrus, inferior temporal gyrus and orbito-frontal cortex, bilaterally.

#### Regression on source localised MEG data

Activity was averaged within each ROI. Then, a linear mixed effects regression model was fit at each time sample separately using the *lme4* package in *R* (Bates et al., 2015). The model included a random slope for trial and subject, and a full random effects structure over items for all of the fixed effects.

#### Connectivity analysis

We used Wiener–Granger causality (G-causality; Granger, 1969; Geweke, 1982) to identify causal connectivity between our different regions of interest in the MEG time series data. This analysis was conducted using the Multivariate Granger Causality Matlab Toolbox [Barnett & Seth, 2014]. The input to this analysis was the time course of activity averaged over all the sources in each Brodmann area of interest. Based on these time series data, we first calculated the Akaike information criteria (AIC; Akaike, 1981) and Bayesian information criteria (BIC; Konishi & Kitagawa, 2008) from our time series data using Morf’s version of the locally weighted regression [Morf, Vieira, Lee & Kailath, 1978]. Then, we fitted a VAR model to our time series data, using the best model (AIC or BIC) order as determined in the previous step, applying an ordinary least squares regression. At this point we checked for possible problems in our data (non-stationarity, colinearity, etc.), and after asserting that our time series fulfilled all the requirements, we continued to calculate the auto-covariance sequence, using the maximum number of auto-covariance lags. Finally, from the sequence of auto-covariance we calculated the time-domain pairwise-conditional Granger causalities in our data. Pairwise significance was corrected using FDR [Benjamini & Hochberg, 1995] at an alpha value of *p* = .01.

## Results

### Behaviour

First, we tested for a behavioural correlate of perceptual adaptation: in this case, faster responses in the picture-matching task as a function of exposure to accented trials. We performed a basic cleaning on the data, removing trials that were responded to incorrectly or faster than 300 ms. Then, we fit a linear mixed effects regression model, including fixed effects for condition (with three levels: baseline, attested, unattested); numerical count of exposures to the speaker; number of elapsed trials; whether they were indicating a match or mis-match with the picture, and speaker identity. Critically, we also included the interaction between condition and number of exposures -- this is the effect we would expect to see if subjects are indeed attuning to the accent. The model also included a random slope for trial and subject, and a full random effects structure over items for all of the fixed effects.

We found that there was no main effect of condition (χ^2^ = 3.23, p = .19),number of exposures (χ^2^ = 0.46, p = .49) or speaker identity (χ^2^ = 0.95, p = .62). There were main effects of number of elapsed trials (χ^2^ = 3.9, p = .048) and match/mismatch response (χ^2^ = 35.34, p < .001). And, critically for our analysis, there was a significant interaction between condition and exposure (χ^2^ = 9.96, p < .01). Average behavioural responses are shown in Figure 2A. Paired t-tests with Holm correction for multiple comparisons revealed a significant difference between baseline and attested (*p* = .046) and between baseline and unattested (*p* = .008) in the first half of the experiment. There was no difference between attested and unattested (*p* = .37), and no differences between any conditions in the second half of the experiment (all *p*’s > .7).

We then fit the same model to the accuracy data. The only significant factor was match/mismatch response, whereby participants were significantly less accurate in mis-match trials (χ^2^ = 147.19, p < .001). Although numerically the baseline trials were more accurately responded to than accented trials 92.92% versus 88.79%, the model did not reveal a significant effect of condition (χ^2^ = 2.51, p = .28) nor an interaction with the amount of exposure (χ^2^ = 1.28, p = .53). There was also no main effect of exposure (χ^2^ = 0.0026, p = .96).

### Neural results

Having identified a behavioral effect of accent adaptation, we then aimed to test whether we could find evidence of perceptual adaptation either in the auditory cortex during sensory processing, or at a higher-level area (i.e. in frontal regions), suggesting a repair process.

To evaluate this, we ran a similar linear mixed effects regression model as the one described above, time-locked to the onset of each critical phoneme (i.e., fixed effects for condition; numerical count of exposures to the speaker; number of elapsed trials; picture match or mis-match, and speaker identity). In addition, we also added phoneme position in the word as a covariate. The model was fit on average responses in transverse temporal gyrus and superior temporal gyrus in the left and right hemispheres separately (our sensory region of interests (ROIs)). And then separately in the orbito-frontal cortex bilaterally (our higher-order ROIs). The model was fit at each millisecond over time, to yield a time-series of model coefficients for each variable of interest. The model coefficients were then submitted to a temporal cluster test in order to estimate during which time window(s) a variable significantly modulated neural responses. The results are shown in Figure 3.

**Figure 3.**
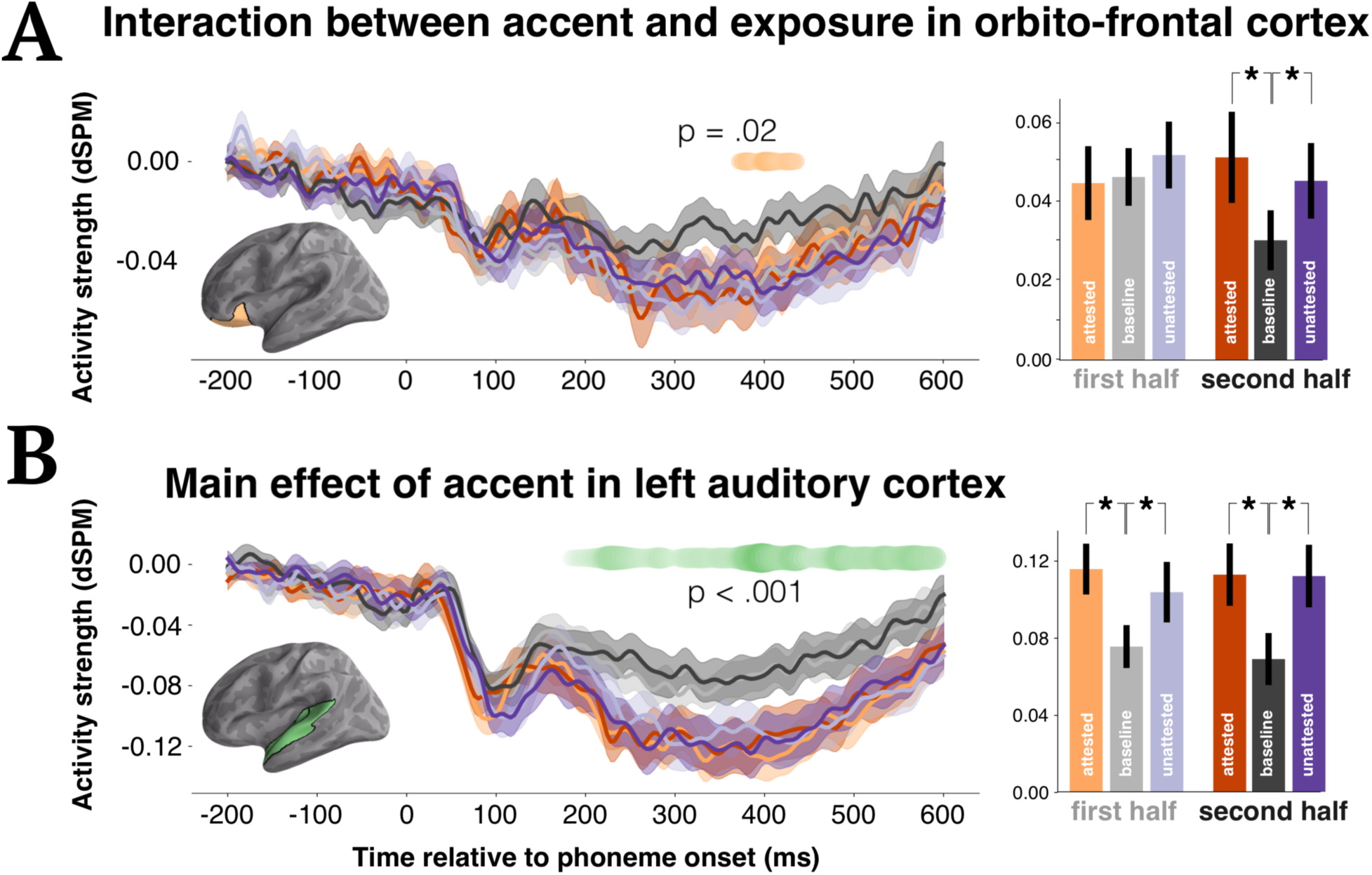
Regression coefficients over time for a multiple regression model with factors: numerical count of exposures to the speaker; number of elapsed trials; picture match or mis-match, speaker identity and condition (baseline, attested, unattested). In each panel, the brain model shows the location of the area where the analysis was condicted. Barplots show activity averaged over the temporal window of significance and over sources in the tested ROI. Panel A) shows an interaction effect whereby prefrontal signals decrease for baseline phonemes over exposure whereas accented speech requires its engagement. Panel B) shows a main effect of accent: regardless of exposure, accented speech always elicits more activity in auditory cortices.

In the left auditory cortex, we found a main effect of condition, such that attested and unattested substitutions consistently elicited more activity as compared to canonical pronunciations, between 188-600 ms (summed uncorrected t-values over time = 51.57; p = .002). There was no interaction with exposure (sum t-value = 6.59), and indeed, numerically the magnitude of the effect remained stable from the first to the second half of the experiment. There were no effects in right auditory cortex.

In the left frontal cortex, there was no main effect of exposure (no temporal clusters formed), but there was a significant interaction between condition (baseline, attested or unattested accent) and exposure from 372-436 ms after phoneme onset (sum t-value = 8.65; *p* = .02). The pattern of the response was such that in the first half of the experiment, all conditions elicited a similar response magnitude, but over exposure, responses to the canonically pronounced phonemes significantly decreased as compared to substituted phonemes. In the right frontal cortex, we instead found a significant effect of condition, responding in a similar pattern as left auditory cortex, also from 376-448 ms (sum t-value = 12.39; p = .033).

We also ran an exploratory analysis by fitting the same model to activity in the inferior frontal gyrus (IFG), superior marginal gyrus (SMG) and middle temporal gyrus (MTG). Overall, we found no effects in IFG or SMG in either hemisphere (cf. Adank et al., 2012). Left MTG showed a main effect of condition in the left hemisphere from between 524-572 ms (sum t-value = 11.53; p = .034). There were no effects in the right middle temporal gyrus.

Overall, the results of the ROI analysis suggest that adaptation to accented speech is primarily supported by processes in frontal cortex. Sensory areas, by comparison, revealed a consistent increase in responses to substituted phonemes. Next we sought to test how information is passed from one region to another.

### Connectivity analysis

Previous theories suggest that listening to accented speech requires the recruitment of additional cognitive processes to facilitate comprehension (Van Engen & Peelle, 2014). Further, recent evidence suggests that not only activity in independent hubs, but also the efficient connection between them, is likely equally as important for successful processing. Based on this, we next sought to test whether the recruitment of additional cognitive processes, in response to accented speech, may be reflected in additional or differential functional connections between brain areas.

To this aim, we ran a Granger causality analysis on our region of interest (ROI) data to determine the connectivity patterns between regions in our data. The complete set of connected is displayed in Fig. 4. Importantly, there are four patterns of results that we want to highlight. First, we found that the bilateral feedforward connection from primary auditory cortex (transverse temporal gyrus) and STG is significant (*p* < .001) for all three experimental conditions. This serves as a sanity check that our analysis is working as intended, given that it is known that these areas interact during normal auditory processing (Pandya, 1995; Kaas et al., 1999). Second, we also find that there are also consistent connections across all conditions between inferior frontal gyrus and primary auditory cortex, bilaterally (*p* < .001) as well as between superior marginal gyrus and superior temporal gyrus, bilaterally (*p* < .001). Third, we only find a significant connection between STG and primary auditory cortex in the baseline condition (*p* < .001), and not in the other conditions. Finally, there is a significant connection between left orbito-frontal cortex and left primary auditory cortex exclusively for the unattested accent condition.

**Figure 4.**
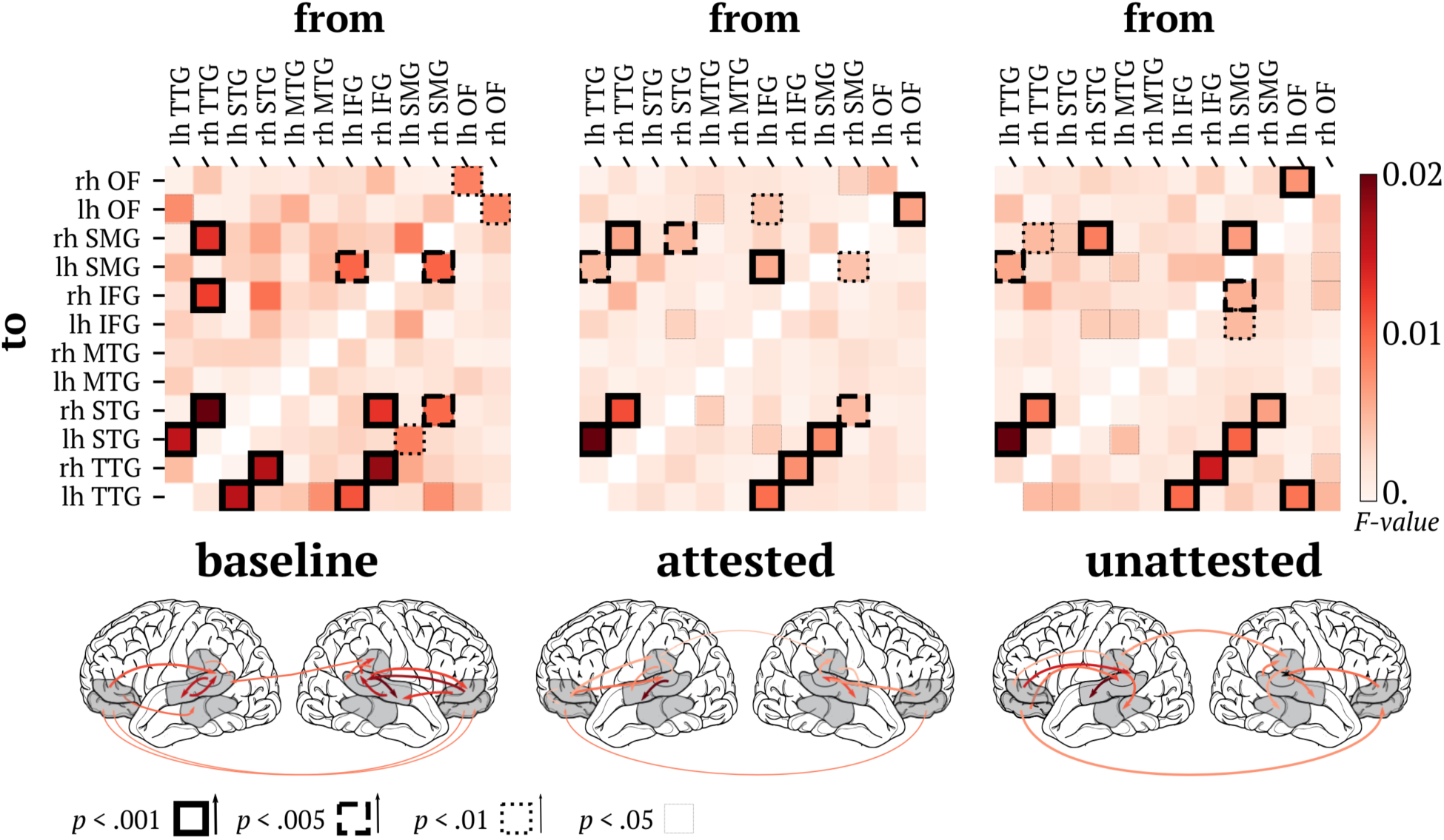
Connectivity patterns across regions bilaterally. In the matrices, the shade of red indicates the Granger causality value of the connections, and the border of the squares the significance level: solid bold line = p < .001; bold dashed line = p < .005; thin dashed line = p < .01; solid thin line = P < .05. The brain models underneath each matrix show the location of the connections with a significance of *p* < .01 or greater. The legend at the bottom indicates that the shade of the line corresponds to strength of the causal connection, using the same colorbar above, and the thickness to statistical significance.

## DISCUSSION

The aim of the present study was to investigate the neural mechanisms supporting perceptual adaptation to accented speech. Previous psychophysics studies have shown that getting used to an accented talker involves recalibrating the mapping from acoustic input to phonetic categories. Here we test whether this recalibration occurs directly at the sensory level (i.e. the listener begins to “hear” the correct phonetic categories directly), or whether it is supported by a higher-order repair mechanism (i.e. the listener initially maps the sounds to the wrong category, and corrects this later). We operationalised this as testing whether the behavioural correlate of accent adaptation manifests itself in the auditory cortices (supporting the sensory hypothesis) or frontal areas (supporting the repair hypothesis).

Overall, our results support the repair hypothesis: the correlate of attunement emerges at a post-perceptual stage, ∼350 ms after phoneme perception in orbito-frontal cortex. However, this correlate did not surface as a decrease in responses to accented speech, which would perhaps have been the most intuitive index of attunement, but rather, as a decrease for unaccented speech that was not mirrored for accents. Further, we found that sensory regions are not modulated by exposure, and consistently elicit stronger responses to accented as compared to canonical pronunciations (180-600 ms). Importantly, the connectivity *between* STG and primary auditory cortex bilaterally is significantly altered by accented speech. The combination of these results has critical implications for speech comprehension specifically, as well as auditory perception more generally.

### Role of frontal cortex

One of the critical findings of the present study is that activity for unaccented speech decreases with exposure in the frontal cortex, whereas responses to accented speech remain relatively elevated throughout. Given the prior association between higher-order cortices and the derivation of conscious percepts (e.g. Del Cul et al., 2009), this response plausibly reflects the coercion of the unanticipated sensory input into a percept that is consistent with the idiosyncrasies of the speaker. This could perhaps unfold under a re-mapping procedure, or a top-down imposition from the lexical context (Samuel, 2001). This therefore suggests that the behavioural speed-up associated with adaptation is a consequence of a stream-lining of this repair process, rather than acting on the sensory input per se.

Since regions of the prefrontal cortex have substantial anatomical connections to auditory areas (Hackett et al., 1999; Romanski et al., 1999), there are anatomical grounds to consider them ideal candidates to modulate the operation of lower-level auditory areas. In fact, lateral portions of the prefrontal cortex have been previously reported to be involved in resolving ambiguity in the identity of speech sounds (e.g., Rogers & Davis, 2012), in speech segmentation (Burton, Small & Blumstein, 2000), and during the comprehension of connected speech (e.g., Humphries et al., 2001; Davis & Johnsrude, 2003; Crinion & Price, 2005; Rodd et al., 2005, 2010; Obleser et al., 2007; Peelle et al., 2010). However, we found that frontal activity only causally modulated auditory cortex activity for unattested accented phonemes. It is possible that the attested condition did not recruit this connection, because subjects were already familiar with these accents. In other words, the substituted phonemes in the attested accent were not “accented enough”, thus failing to recruit the full repair mechanism.

### Role of auditory cortex

The second critical result of this study is that accented speech elicited consistently increased activity in auditory cortex. Unlike activity in frontal areas, these responses *did not wane* as a function of exposure. This pattern of results is interpretable under a predictive coding framework – the idea that the brain continuously generates predictions about upcoming sensory input, and elicits increased responses when the expected and received inputs do not align. Here, of course, accented speech can be construed as the less expected input. Importantly, even though a listener may have the conscious experience of being able to better predict the pronunication of a speaker with time, this is not reflected in the update of prediction signals in auditory cortex.

These predictive error signals have been found both during lexical (Gagnepain et al., 2012) and non-lexical (Dürschmid et al., 2016) contexts in audition, as well as in perception during object vision (Egner, Monti, & Summerfield, 2010) and multimodal integration (Arnal, Wyart, & Giraud, 2011; von Kriegstein & Giraud, 2006), positing them as a general principle under which the sensory brain operates, as opposed to them being a speech-specific mechanism. In the case of speech, this response has been linked to modulation of activity around 200-300 ms in transverse temporal gyrus and superior temporal gyrus (Gagnepain et al., 2012; Ettinger et al., 2014; Gwilliams & Marantz, 2015; Gwilliams et al., 2018). The timing (200-600 ms) and location (STG) of our observed effects are thus consistent with previous studies and posits prediction error signals as a plausible source of hightened response to accented speech. Importantly, there were no effects of accent at the auditory M100 component. This is likely due to the fact that this response reflects bottom-up processing, without considering the global lexical content of the word, and at the lowest-level, all the phonemes in our stimuli were well-formed - even the accented ones.

Finally, we found that both types of accented speech disrupted existing connections between STG and primary auditory cortex, as evidenced by our connectivity analysis. Although it is conjecture at this point, it is possible that the additional connections recruited for unaccented speech result in more efficient processing. And, further, if local connections are in place within sensory areas, there is less need to recruit additional external regions during processing. However, future investigation is necessary to substantiate this claim.

### Implications for models of effortful listening

The general model of effortful listening proposed by Van Engen and Peelle (2014) suggests that the comprehension of accented speech is cognitively effortful because there is a mismatch between the expectations of the listener and the received signals. Further, they intuitively suggest that getting used to a given accent should automatically change the expectations of the listeners, hence reducing the mismatch between expectations and input. This should in turn lower the demand for compensatory signals and associated cognitive effort.

Our experiment did not straightforwardly match these predictions: the signals to accented speech in auditory cortex remained constant throughout, and there was no difference between an accent that participants had arguably heard before versus not. This opens two possibilities. The first is that the effortful listening model is innaccurate in assuming an effect of social expectation in auditory perception. At the lowest level, phonemes will inevitably be categorized based on the physical properties of the signal, and not based on the intended phoneme of the speaker. Thus, there is no escaping automatic predictions, and subsequent prediction error signals, in auditory cortex.

The second possibility is that there is a dissociation between behavioral and neural adaptation timelines. That is, even if behaviorally adaptation occurs quickly, the underlying neural processes will adapt at a slower pace and will necessitate two stages. Although our study did not contain enough exposure to dissociate between these two stages, theoretically, this is how such a mechanism would work. First, after a short exposure to an accent, listeners will succeed at fully understanding accented speech, as shown by behavioral accuracy measures. However, achieving this intelligibility will require additional cognitive effort, as reflected in i) elevated difficulty in understanding (Munro & Derwing, 1995a; Schmid & Yeni-Komshian, 1999) and ii) slower processing for accented speech (Munro & Derwing, 1995b; Floccia et al., 2009), and it will rely on high prediction error *correction* signals. Further down the line, the effort required to comprehend will reduce: we predict that at this stage there will be a decrease in the error prediction signal, and a reestablishment in the functional connections between STG and primary auditory cortex, leading to a pattern of activity more similar to that of the perception of canonical speech. Because our experiment gave people relatively little exposure, our results do not adjudicate between these two possibilities.

## CONCLUSION

In all, this paper constitutes the first characterization of the neural markers of auditory adaptation to foreign accents during online perception of speech sounds. These findings contribute a critical piece to our hitherto poor understanding of how the brain solves this puzzle, which was reduced to behavioral descriptions of this phenomenon, and anatomical structural differences for better accent perceivers (Golestani, Paus & Zatorre, 2002). We unveil that understanding foreign accents relies on prediction error signals and propose that this only represents the first stage on the road to attunement, which will lead to a lower level adaptation of sensory classification by hypothesis.

## Acknowledgements

This work was funded by the Abu Dhabi Institute grant G1001 (A.M. & L.P.) and the Dingwall Foundation (E.B.E & L.G). We thank Lena Warnke for her assistance in stimuli selection.

